# Moremi Bio Agent: Using *Neisseria meningitidis* Reference Data For The Double Blinded Validation of A General Purpose Biology-Trained Reasoning Model for Pathogen and Antigen Discovery

**DOI:** 10.64898/2025.12.17.694980

**Authors:** Gertrude Hattoh, Solomon Eshun, Nyarko Prince Ofori, Mohammed Nuruddin Alhassan, Fadil A Bidmos, Darlington Akogo

## Abstract

Antibodies serve as vital diagnostic and therapeutic agents due to their exceptional specificity toward antigenic targets. Mapping antibody–antigen interactions is essential for understanding immune responses and developing vaccines or biologics. Traditional antigen identification relies on labor-intensive wet-lab techniques such as phage display, peptide microarrays, and ELISA, while computational methods employ sequence alignment, epitope mapping, and structure prediction. Despite progress, to our knowledge, no existing AI framework has demonstrated the ability to blindly inference—predicting an antibody’s antigen target solely from its amino acid sequence without prior biological context. This research employed Moremi Bio Nano, a general agentic reasoning large language model (LLM), to infer the antigen and pathogen targets of an anonymized monoclonal antibody sequence from Imperial College London. The model received only the VH and VL chain sequences and autonomously hypothesized, ranked, and validated probable targets. Of ten independent inference tests, four were completed successfully, with three correctly identifying the experimentally validated antigen and pathogen; with SARS-CoV-2 Spike RBD, *Neisseria meningitidis* fHbp v1.1, and SARS-CoV Spike emerging as the top 3-ranked candidates across both reranking strategies. Validation of the model’s predictions with experimental wet-lab data confirmed its capacity for correct antigen inference, marking Moremi Bio Nano as a first-of-its-kind AI system demonstrating reasoning-driven antigen discover; complementing experimental immunology and advancing automated biological inference.

## 1. INTRODUCTION

Antibodies constitute one of the most versatile and indispensable molecular tools in modern medicine, diagnostics, and therapeutic design [1]. Whether produced naturally by plasma cells as part of the humoral adaptive immune response 1, or recombinantly or by hybridoma/ plasma cell technologies, monoclonal antibodies (mAbs) possess exceptional specificity and affinity toward unique antigenic epitopes 2. These properties have established mAbs as critical components in immunotherapy, targeted drug delivery, and disease diagnosis.

Clinically approved mAbs such as Trastuzumab (HER2-positive breast cancer), Adalimumab (TNF-*⍺* inhibition in autoimmune disorders) and Palivizumab (RSV prophylaxis) exemplify their broad therapeutic relevance. Beyond therapeutic applications, mAbs are essential in diagnostic assays—including Enzyme Linked ImmunoSorbent Assay (ELISA), immunoprecipitation, and rapid immunochromatographic tests—where their precision in antigen detection (identification and characterization) enables sensitive biomarker identification [2].

The identification and characterization of antigenic targets form the cornerstone of immunological research and antibody-based therapeutic discovery. Determining the specific immunogenic component and/or epitope recognized by an antibody’s paratope provides insight into binding affinity, neutralization potential, and off-target liabilities. In addition, it powers the identification of novel therapeutics for disease states, and pushes the frontier for easier therapeutic design. However, conventional antigen detection methods—such as ELISA [3], mass spectrometry, phage display, or alanine-scanning mutagenesis, while effective, are often laborious, time-consuming, and resource-intensive. These limitations become especially pronounced in cases involving novel or poorly characterized pathogens, where reference data and clinical precedents are scarce.

Rapid and accurate antigen–antibody mapping therefore represents a critical unmet need, as it enables faster pathogen characterization, accelerates therapeutic development, and supports diagnostic precision during emerging infectious disease outbreaks. The challenge of antigen–antibody mapping remains a bottleneck: conventional laboratory methods such as ELISA, mass spectrometry, phage/yeast display or alanine-scanning mutagenesis frequently demand substantial time and resources, particularly when dealing with novel or poorly characterised pathogens.

Computational strategies have significantly improved antigen discovery workflows. Techniques such as sequence alignment, structural modeling, and supervised machine learning trained on antibody–antigen pairs [4] facilitate high-throughput prioritization of potential targets. Nevertheless, these models typically depend on well-annotated training datasets or predefined antigen families. Their generalizability diminishes when confronted with novel antigens or antibodies derived from unrepresented proteomes, thus restricting their applicability in real-world, data-sparse discovery contexts [5].

Recent advances in artificial intelligence (AI) 3 have transformed drug discovery, structural biology, and molecular design. Systems such as AlphaFold, ESMFold, and related protein language models have demonstrated remarkable accuracy in predicting protein folding and structure [6][7]. Recent surveys on existing AI and LLMs reveal that no current AI system can reliably infer a novel antibody’s antigen target or associated disease state de novo – that is, directly from antibody sequence data without prior antigen information. Although numerous models have advanced antibody–antigen interaction prediction, [8][9][10][11], these approaches typically assume prior knowledge of the antigen sequence or structure and function primarily as discriminative classifiers or structure-based docking engines rather than as reasoning systems capable of reverse antigen inference – i.e infer antigen target de novo from antibody sequence alone.

Notably, platforms such as AI-Immunology^TM^ (Evaxion Biotech) represent a significant advance in forward antigen discovery, particularly for vaccine development, by leveraging machine-learning models (e.g., EDEN^TM^) trained on curated immunogenicity datasets to rank pathogen-derived antigens likely to elicit protective immune responses [12]. While highly effective for proteome-scale antigen prioritization and vaccine design, AI-Immunology^TM^ is not designed to infer the antigenic target of an existing monoclonal antibody from sequence alone, nor to reason backward from antibody paratope features to pathogen identity. Consequently, its predictive paradigm differs fundamentally from the reverse-inference task addressed in this study.

A further limitation common to many prior models—including both interaction predictors and antigen-ranking systems—is training-data bias toward well-studied antigens (e.g., viral surface glycoproteins) and human therapeutic antibodies, which constrains generalization to rare pathogens,unusual antibody repertoires, or blinded inference settings [13].

Despite remarkable advances in artificial intelligence (AI) and LLMs applied to biological sequences, to our knowledge no publicly reported system has been validated to infer a novel antibody’s precise antigenic target and associated disease state or pathogen ab initio—that is, without prior knowledge, templates, or contextual data [14]. This limitation to our knowledge constrains computational immunology and underscores the need for reasoning-based, context-independent systems that can bridge the divide between molecular recognition and immunodiagnostic reasoning.

To address this gap, we present Moremi Bio Nano (a new version of Moremi Bio Agent [15][16] an agentic, general purpose reasoning LLM for biomedical and drug discovery research, that has shown the potential for sequence-agnostic antigen discovery (I.e without contextual data or prior model training specifically on this task). Although Moremi Bio Nano has implications for both therapeutic antibody design and diagnostics, in this study we emphasise its utility as a discovery tool for early-stage antigen target identification and diagnostic marker mapping. We specifically evaluate its ability to infer disease state(s), pathogen(s), and precise antigenic targets directly from the amino acid sequences of novel monoclonal antibodies alone. The model was tested in a blinded setting using an antibody sequence cloned from a healthy individual and shared by the Imperial College Group. The goal was to determine whether Moremi Bio Nano could independently identify the same pathogen and antigen experimentally validated by wet-lab assays. By narrowing the predictions to the top five probable antigenic targets per antibody, this study seeks to:

- Validate the reasoning capacity of Moremi Bio Nano in antigen inference without auxiliary datasets;
- Compare its predictive outcomes with laboratory-confirmed results; and
- Assess its suitability as an *in silico* tool for early-stage drug discovery, immunodiagnostics, and immunogenic target validation.

Here we propose a new paradigm in antibody-based antigen discovery: an AI-driven framework capable of reasoning directly from antibody primary sequence without reliance on prior antigen annotation. If validated, Moremi Bio Nano could markedly accelerate immunogen discovery pipelines, reduce experimental overhead, and enhance preparedness for emerging pathogens by enabling de novo antigen prediction at scale.

### 1.1 Moremi Bio Nano

Moremi Bio Nano,( I.e Moremi Bio Agent 4 v2) is a general purpose agentic and reasoning LLM optimized for biological, biomedical and drug discovery research. The model couples a transformer-based reasoning model with explicit agentic orchestration layers that dispatch specialized bioinformatics and computational toolchains (3D structure generation, docking, ADMET prediction, molecular simulations) and query external curated resources (RCSB PDB, IEDB, PubChem, Enamine DB and in-house micro-DB, derived from PDB databank) for evidence-grounded inference. It implements task-based reasoning chains and multi-mode inference, a staged pipeline that alternates hypothesis generation, symbolic/knowledge retrieval, tool execution, and iterative validation allowing variable computational investment proportional to task complexity.

Importantly, the model was not trained on antigen–antibody discovery tasks, nor was bacterial antigen discovery included in its pretraining corpus. The emergence of accurate antigen inference in this study therefore reflects a novel, untrained capability arising from its reasoning framework rather than dataset memorization, underscoring the scientific significance of the results. This task presents an emerging capability, such emergent behavior suggests that advanced reasoning-centric LLMs may be approaching a threshold where they can perform foundational biological inference traditionally requiring experimental pipelines.

**Figure 1:**
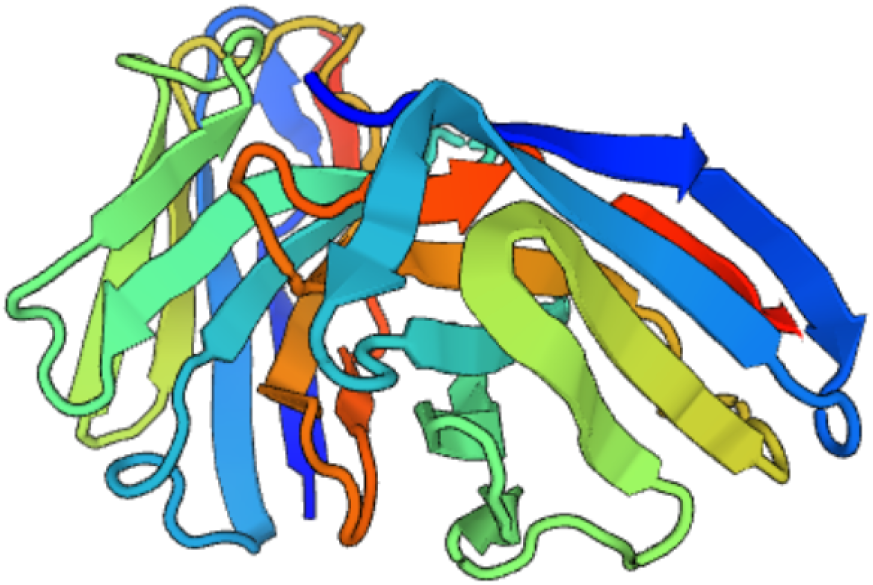
Structure of Antibody

The architecture therefore unifies declarative knowledge retrieval, procedural simulation, and reflective decision-making for reproducible in silico candidate discovery and prioritization. Moremi Bio Nano inherits the full computational toolchain of its predecessor (Moremi Bio Agent) and extends it with additional capabilities, expanding both the breadth and depth of its analytical repertoire. A summarized list of previously available tools is provided 5, upon which the current system builds with further integrations and enhanced reasoning-driven orchestration [Read from the supplementary data 5, for sample reasoning chain of Moremi Bio Nano].

## 2. METHODOLOGY

### 2.1 Input Data and Sequence Information

VH and VL (*κ*) sequences of a fully human monoclonal antibody (hmAb VFB-8D5), cloned from a healthy donor prior to the COVID-19 pandemic, were employed in this study. Prior characterisation has been reported [17]; however, the identity of the cognate antigen determined by protein-array screening remains unpublished. The cognate antigen identity and clinical metadata (volunteer history and prior immunization) were blinded between experimental and computational analyses.

### 2.2 Experimental Objective

The objective was to evaluate whether Moremi Bio Nano, an agentic reasoning large language model (LLM), could accurately infer:

- The disease state and pathogen associated with the antibody sequence.
- The precise antigenic target of the antibody, solely from its primary amino acid sequence, without contextual information or prior training exposure to the sequence.

The wet-lab reference (true pathogen and antigen) was known only to the Imperial College team and was withheld from all computational researchers at MinoHealth AI Labs to maintain experimental blindness.

### 2.3 Model testing Framework

The model was tested yielding three successful inferences, each conducted in a separate, isolated chat session to prevent contextual contamination across runs. Within each session, the model was prompted with the same antibody sequence and instructed to:

- Identify potential pathogen(s) and associated disease(s) targeted by the antibody.
- Infer the most probable antigen(s) recognized by the antibody.
- Rank the top five predicted targets based on confidence scores derived from reasoning outputs and internal evaluation metrics.

Each session consisted of ≈5–6 interaction cycles between the model and user prompts. User input was minimized and limited to non-directive confirmations (e.g., ‘proceed’ or ‘continue’) to preserve autonomy in model reasoning yet still with human supervision. The model determined its own analytical pipeline—selecting computational tools, ordering their use, and formulating hypotheses independently, reflecting its autonomy in reasoning yet with human supervision.

### 2.4 Analytical Workflow

Across all inference sessions, Moremi Bio Nano autonomously selected and executed tools from its integrated suite, spanning multiple computational domains: Sequence-based analyses, epitope prediction and mapping, structural inference, binding and affinity estimation and database integration which includes in-depth literature, along with structural database cross-referencing. The model generated ranked hypotheses supported by internal reasoning and cross-validation across tools. In Strategy Two, which incorporated detailed tool outputs and experimental logs from each test, the model processed up to 28,243 tokens in a single query (Table 1), demonstrating its capacity to handle extremely large, complex datasets while maintaining coherent multi-step reasoning. Across the completed runs, Moremi Bio Nano used 35 minutes (minimum inference time) to 131 minutes (maximum inference time) per session. This extended time expenditure is consistent with agentic-model behavior, in which prolonged deliberation reflects deeper multi-step reasoning, tool orchestration, and iterative hypothesis refinement. The average inference duration (≈ 1ℎ*r*23 minutes per session) reflects intensive computational reasoning required to infer immunological targets from sequence alone, a task traditionally considered refractory to automation and dependent on wet-lab validation.

**Table 1:** Comparison of Strategy One and Strategy Two.

| Strategy | Description |
| --- | --- |
| Strategy One | The summary result tables from each of the four completed tests were consolidated and presented to Moremi Bio Nano in a single query. The model was tasked with reranking all antigens collectively based on their relative confidence, identity, and inferred biological plausibility. |
| Strategy Two | Detailed tool outputs and experimental logs from each test—together with the corresponding summary tables—were provided to Moremi Bio Nano for reranking (Top to 28243 tokens used). This enabled the model to incorporate both intermediate computational evidence and final task-level outcomes during prioritization. |

### 2.5 Completion Evaluation Criteria and Validation Benchmark

Four (4) were successfully completed and qualified for inclusion in the final analysis. A session was considered complete when the model:

- Produced a full ranked list of five predicted antigen/pathogen pairs.
- Recommended a validation workflow (e.g., ELISA, surface plasmon resonance, or structure refinement) to confirm its predictions experimentally.

The predictions generated by Moremi Bio Nano were subsequently compared against experimentally verified wet-lab data from the Imperial College research group. Agreement between the top-ranked model prediction and the laboratory-confirmed antigen served as the primary validation metric, thereby assessing the accuracy and reasoning capacity of the model for de novo antigen discovery.

### 2.6 Re-ranking of 4 successfully Completed sessions

To identify the most probable antigen targets across all four successful tests, two independent reranking strategies were implemented. In both approaches, the agentic LLM Moremi Bio Nano served as the evaluator, autonomously reanalyzing and ranking the antigen candidates. The reranking processes were conducted in separate experimental sessions to ensure independence and reproducibility.

Notably, both reranking strategies yielded identical Top-4 ranked pathogens and antigens, indicating consistency and robustness in the model’s inference process [See Appendix B].

## 3. RESULTS

### 3.1 Overview of test sessions

Tests were conducted using the same query. Out of these, some were incomplete due to tool execution errors or unresponsive user input. All completed tests were included in the final re-ranking analysis.

In all completed sessions, Moremi Bio Nano reasoned and employed its agentic system to generate ranked predictions of five potential antigen–pathogen pairs per test. In three of the four completed tests, the model successfully identified the precise wet-lab validated antigen and its corresponding pathogen, achieving full inference accuracy. Notably, in each of these successful tests, the correct pathogen–antigen pair consistently appeared at Rank 2 in the model’s prioritized output list.

Across the four successful tests, the model autonomously selected and executed a total of 47 tool calls. Each successful run employed an average of 7.5 ± 2.38 tool executions, with 4.25 ± 1.50 tool attempts failing due to parameter mismatches or timeouts [See Appendix A]. This demonstrates robust but imperfect orchestration of *in silico* analytical tools within an agentic framework.

### 3.2 Re-ranking and Consensus Analysis

From the re-ranking test, both strategies produced identical Top-4 antigen–pathogen rankings [See Appendix B]. The average confidence bands were taken, and below is a graph of model inference accuracy and targets from the reranking test.

### 3.3 Validation Against Wet-Lab Data

The summary table of test 1 and 8 was submitted to the Imperial College London research group for validation. Cross-referencing with wet-lab data confirmed that the **second-ranked antigen–pathogen pair (*Neisseria meningitidis fHbp* v1.1)** was indeed the correct and experimentally validated target of the sequenced monoclonal antibody. This outcome establishes the **first recorded successful blind inference** of a true antigen–pathogen target **from antibody amino acid sequence alone** by an LLM framework, thus validating **Moremi Bio Nano** as a credible *in silico* reasoning system for biomedical research.

**Figure 2:**
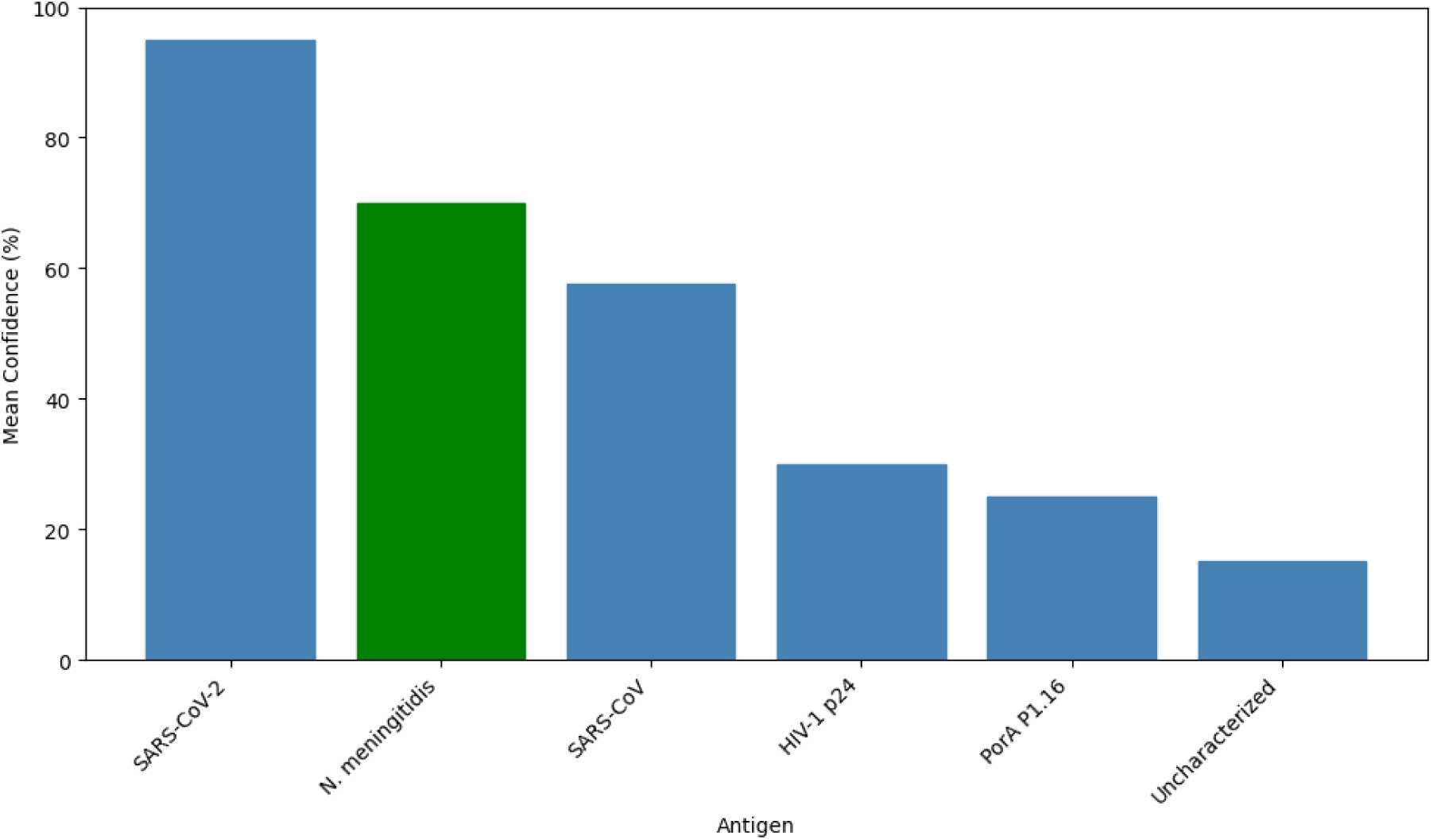
*In silico* inference of antibody-antigen specificity. Mean confidence scores (%) for the top-ranked antigen candidates predicted from a blind antibody sequence input. The green bar denotes the experimentally validated target (*Neisseria meningitidis* fHbp v1.1), confirming the model’s capacity for accurate target identification among high-confidence predictions.

## 4. DISCUSSION

Identifying the precise antigenic target of a monoclonal antibody solely from its amino acid sequence remains one of the most complex challenges in immunoinformatics. Conventional antigen discovery methods, such as phage display, ELISA and X-ray crystallography, require extensive wet-lab experimentation and are resource-intensive. Computational approaches, while more scalable, depend largely on existing antigen–antibody complex databases or known sequence similarities, which restricts their ability to generalize to novel or uncharacterized antibodies.

In this study we evaluated Moremi Bio Nano in a fully blinded experiment comprising N =1 novel mAb sequence, of which it had experimentally validated antigen/disease target from Imperial college Group. By treating the antibody sequence as input and leveraging the agentic reasoning nature of the model, Moremi Bio Nano can hypothesise plausible antigen contexts (e.g., pathogen) even in absent explicit antigen training pairs, thereby enabling generalisation to novel or under-represented proteomes. In this work, we define an antigenic target as a specific pathogen-derived protein to which the antibody binds; for disease mapping we infer the likely pathogen and hence associated disease state. Metrics included top-5 accuracy, and specificity across disease-pathogen-antigen tiers. The wet lab validated results for the antibody used in this test was *Neisseria meningitidis* fHbp v1.1, which appeared in 3/4 successful tests and always placed at rank 2.

SARS-CoV-2 Spike receptor-binding domain (RBD) consistently occurred in all four of the successful inference sessions and was placed top-rank in all tests. Although the wet-lab validated antigen is fHbp, the persistent prioritization of Spike RBD by the model raises a concern on the probability of the model’s bias towards immunodominant, well-represented antigens. RBD is a known immunodominant epitope: studies show that over 90% of neutralizing antibodies in SARS-CoV-2 immune sera target the RBD region of the spike protein [18].

The recurrent identification of the SARS-CoV-2 Spike RBD as the top-ranked antigen across all successful inference sessions is a striking pattern given that the antibody in this study was collected and cloned from donor in summer 2019, months before the recognized onset of COVID-19 pandemic. This chronology excludes the possibility of classical immune imprinting by SARS-CoV-2 and therefore challenges the assumption that the sequence features could arise from direct viral exposure. Nonetheless, the model’s consistent prioritization of Spike RBD suggests that the antibody sequence contains structural or paratope-level features that resemble motifs enriched in public anti-RBD clonotypes which was frequently highlighted in observations such as CDR3 similarity to published anti-RBD clonotypes. It is notable that emerging retrospective analyses have detected SARS-CoV-2–reactive antibodies in samples from as early as March 2019, raising the possibility that antigenically similar betacoronaviruses may have been circulating earlier than clinically recognized and could, in principle, shape baseline immunological landscapes [19].

While the pre-pandemic sample date in this study argues against direct exposure to SARS-CoV-2 itself, the persistent computational signal may still reflect deeper biological phenomena such as epitope mimicry, germline-encoded cross-reactivity, or structural convergence between fHbp-targeting antibodies and RBD-binding public clonotypes. Alternatively, if an antibody of this study had arisen after the pandemic, one could plausibly posit an evolutionary path in which an early SARS-CoV-2–directed precursor later diversified and matured toward fHbp; yet, this is not the case for this study. We therefore plan to synthesize the Spike RBD protein in the lab and experimentally evaluate binding and functional activity. This follow-up will clarify whether the model’s persistent identification of Spike RBD reflects a genuine underlying biological process and intent to validate the model’s suggestion.

**Figure 3:**
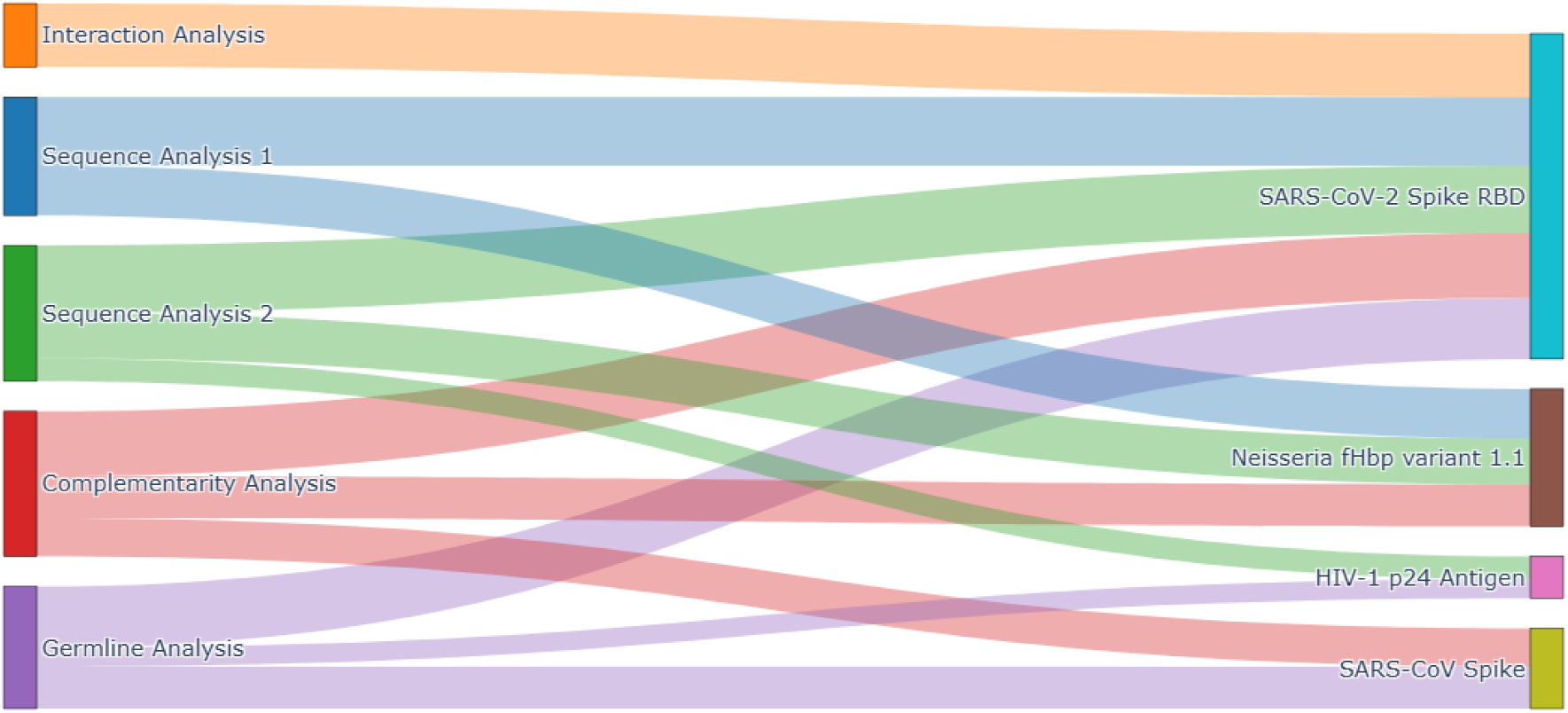
Integrative Evidence Flow Mapping for Antigen Prioritization by Moremi Bio Nano Across Successful Inference Tests

This Sankey diagram illustrates the relative contribution of diverse immunoinformatics evidence streams —Sequence analysis, germline analysis, complementarity analysis, and interactions analysis —toward the identification and prioritization of likely antigenic targets inferred by Moremi Bio Nano. The weighted flows quantify the inferred confidence and multi-layered convergence of antibody–antigen associations across distinct analysis modules. This representation provides a transparent overview of how heterogeneous molecular evidence streams collectively inform antigen ranking (SARS-CoV-2 Spike RBD as rank 1) within the reasoning-based antibody target inference pipeline.

Our results indicate that Moremi Bio Nano was able to reason over multiple layers of evidence and propose ranked antigen–pathogen lists, with the validated wetlab target consistently appearing at Rank 2 in the majority successful runs. This suggests the model has learnt a heuristic capable of prioritizing biologically plausible antigen candidates even in the absence of clinical metadata or immunization history. The consistency between two independent reranking strategies further underscores the model’s internal reasoning stability.

Moreover, the study included one novel antibody sequence (N=1) and focused on healthy-donor derived mAbs; it remains unclear how performance scales with hypermutated antibodies, uncommon pathogens, or highly glycosylated antigens. Expanding testing across diverse experimental designs is therefore a critical next step to establish reliability and generalizability before considering translational or diagnostic applications.

### 4.1 LIMITATIONS

We emphasise that this study focuses on inference from sequences of an isolated antibody (i.e., with only Variable heavy and light chains, yet without extensive metadata) and that successful inference in this reduced-context regime remains highly challenging. Identifying the precise antigen targeted by an antibody solely from its amino acid sequence remains an exceptionally complex task due to the vast diversity of the human immune repertoire, the context-dependent nature of antibody–antigen interactions, and interindividual variability in immune responses that influence somatic hypermutation and epitope recognition [20][21][22]. We acknowledge that antibody binding specificity arises from 3D complementarity, somatic hypermutations, and often complex post-translational modifications; hence sequence-only inference remains inherently probabilistic rather than deterministic, and our model is intended for hypothesis generation and validation (which intend require validation of wet lab results) rather than definitive assignment.

## 5. CONCLUSIONS

This study provides proof-of-concept that an agentic LLM can infer antigen–pathogen targets from novel antibody sequences without prior contextual data, thereby bridging a critical gap in immunoinformatics. With further refinement, such systems may transform early-stage drug discovery, immunodiagnostic development, and rapid therapeutic responsiveness for emerging pathogens.

## APPENDICES

## A. TOOL CALLS

This table summarizes the number of successful and failed tool calls executed by Moremi Bio Nano across the four completed inference sessions (Tests 1, 8, 9, and 10).

**Table 2:**
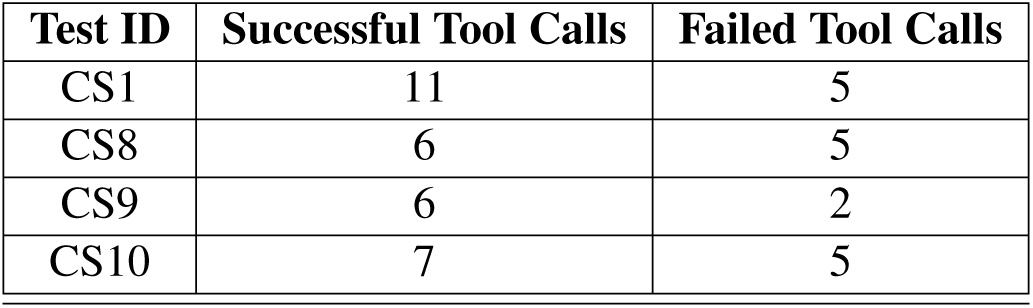
Tool usage outcomes across independent inference tests.

## B. RESULTS FROM RERANKING OF 4 SUCCESSFULLY COMPLETED SESSIONS

**Table 3:** Re-Ranked Antigen Targets for the Novel Monoclonal Antibody (hmAb-VH/hmAb-Vk) from ranking test using results + tools response logs (Strategy 2). This shows the precise associated disease state and predicted antigen target of each rank.

| Rank | Antigen Target | Disease / Pathogen | Confidence Band |
| --- | --- | --- | --- |
| 1 | SARS-CoV-2 Spike RBD | SARS-CoV-2 (COVID-19) | High (95%) |
| 2 | fHbp variant 1.1 | *Neisseria meningitidis* | Medium (70%) |
| 3 | SARS-CoV Spike | SARS-CoV (SARS) | Medium (65%) |
| 4 | HIV-1 p24 Antigen | HIV-1 | Low (30%) |
| 5 | Uncharacterized Ig Variable Regions | N/A | Low (< 15%) |

## C. CONSOLIDATED RERANKING RESULTS

**Table 4:**
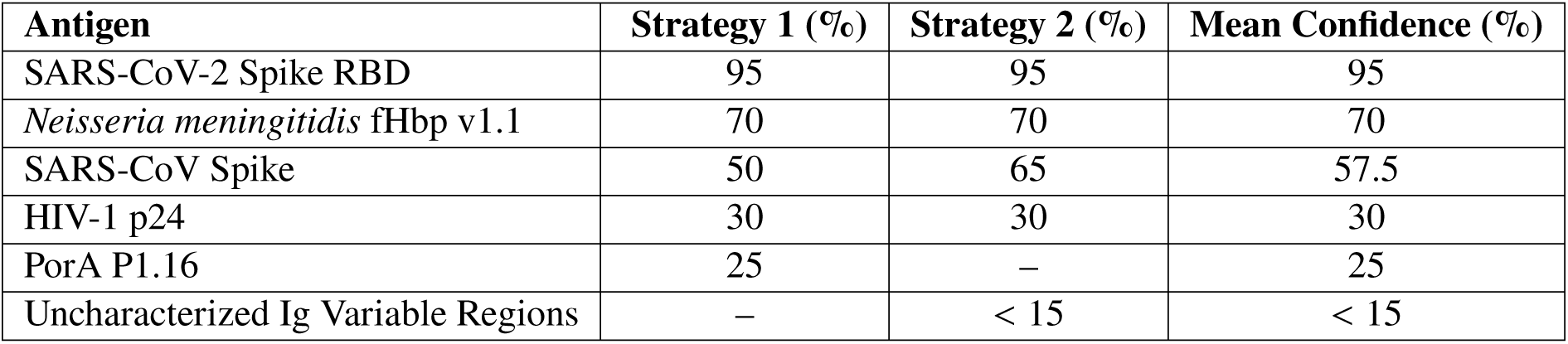
Consolidated reranking results of all antigen–pathogen candidates across completed test sessions.

| Antigen | Strategy 1 (%) | Strategy 2 (%) | Mean Confidence (%) |
| --- | --- | --- | --- |
| SARS-CoV-2 Spike RBD | 95 | 95 | 95 |
| *Neisseria meningitidis* fHbp v1.1 | 70 | 70 | 70 |
| SARS-CoV Spike | 50 | 65 | 57.5 |
| HIV-1 p24 | 30 | 30 | 30 |
| PorA P1.16 | 25 | – | 25 |
| Uncharacterized Ig Variable Regions | – | < 15 | < 15 |

## SUPPLEMENTARY DATA

### Illustrative Reasoning and Tool-Orchestration Snapshots

To provide transparency into the internal operation of the agentic AI framework, we include a limited set of illustrative snapshots depicting representative reasoning chains and tool-orchestration behaviors generated by the model during an independent, non–antigen-discovery use case. These examples are presented solely to demonstrate how the model decomposes complex scientific queries, sequences intermediate reasoning steps, and dynamically invokes external computational tools, rather than to support or influence the biological conclusions of the present study.

The showcased session is not related to the antigen identification task analyzed in the main manuscript, and no biological inference from these examples is used in model evaluation or interpretation. Substantial portions of the reasoning traces have been intentionally obscured to protect proprietary model logic and prevent over-interpretation of incomplete intermediate states, while preserving sufficient structure to illustrate the model’s methodological workflow.

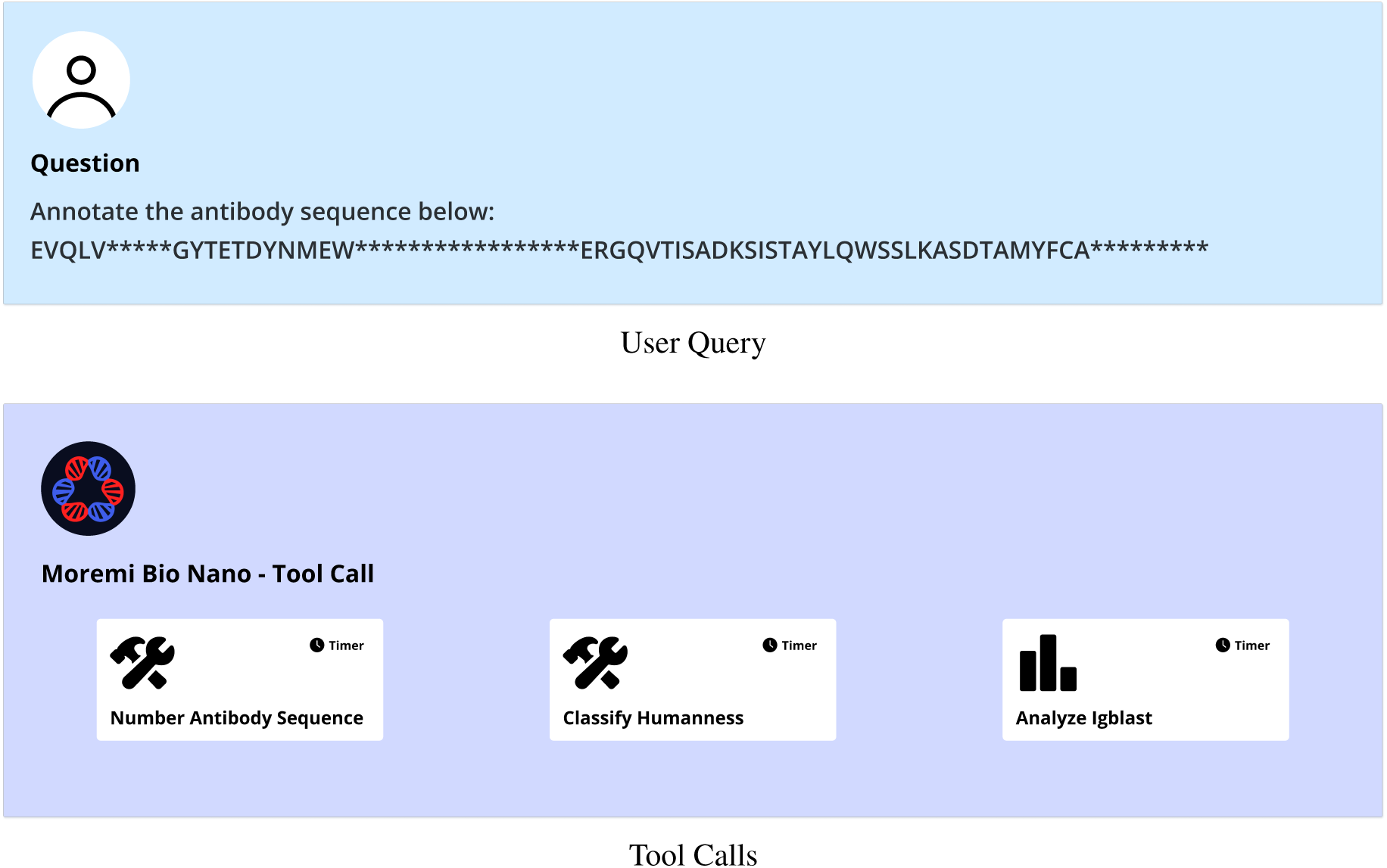

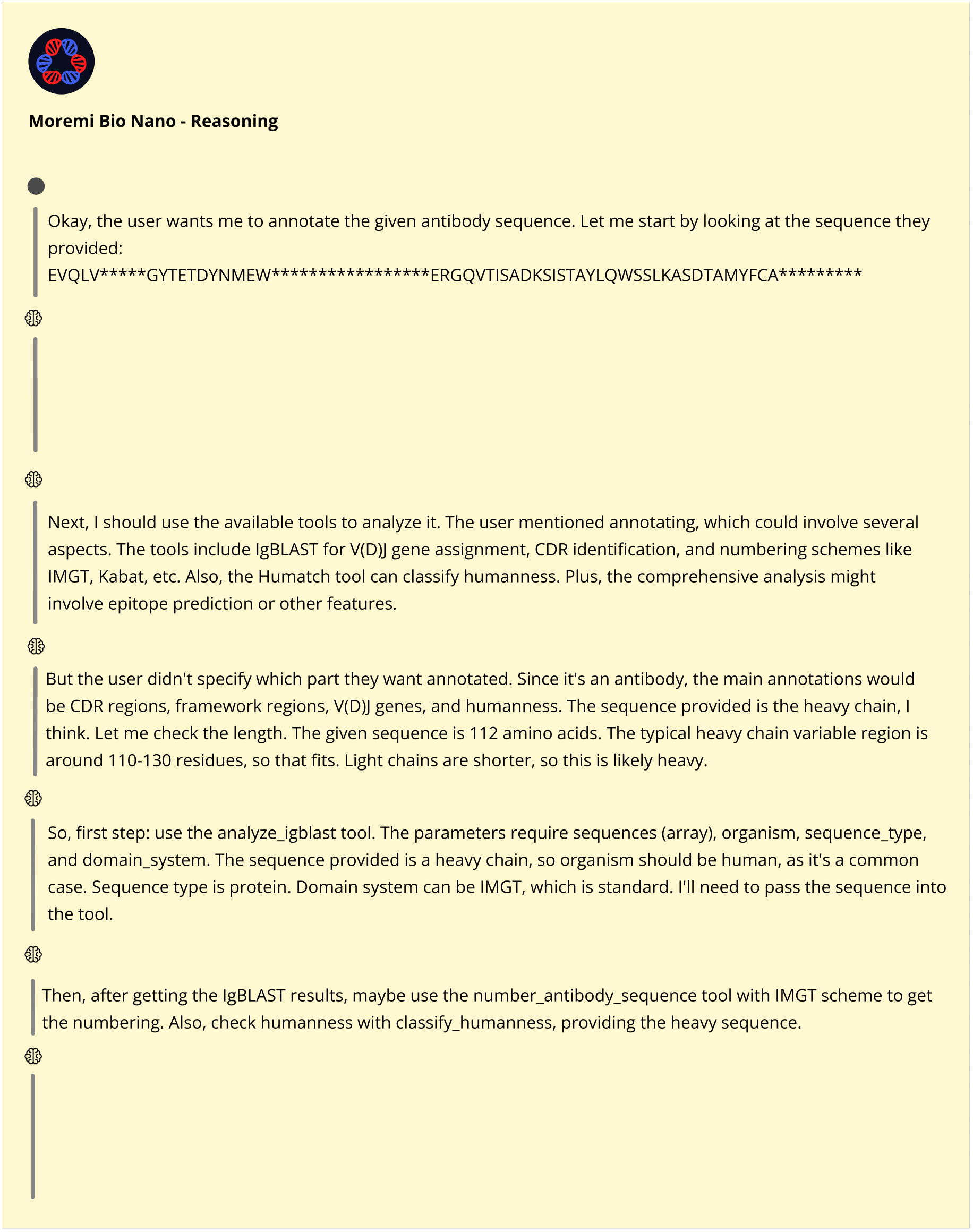

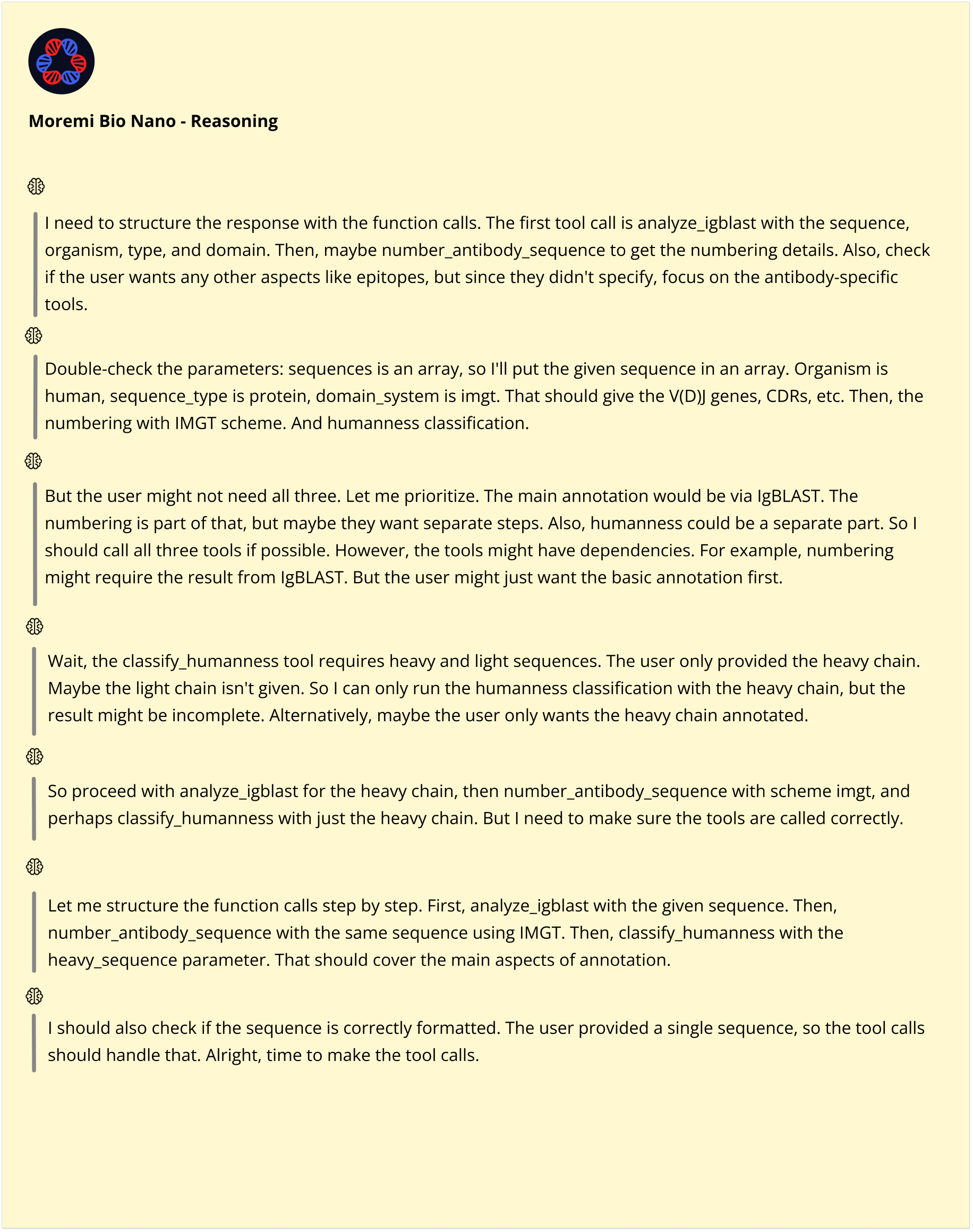

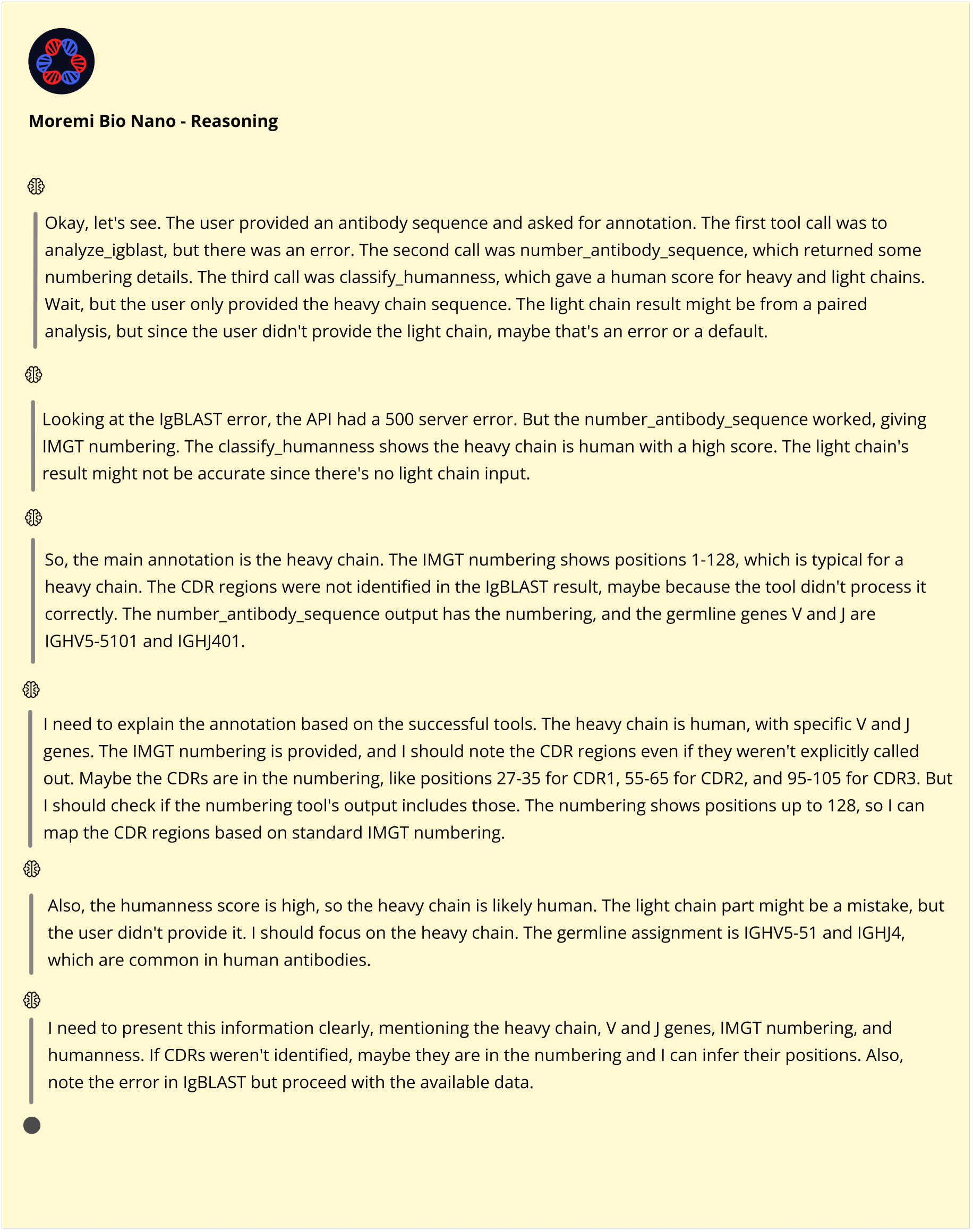

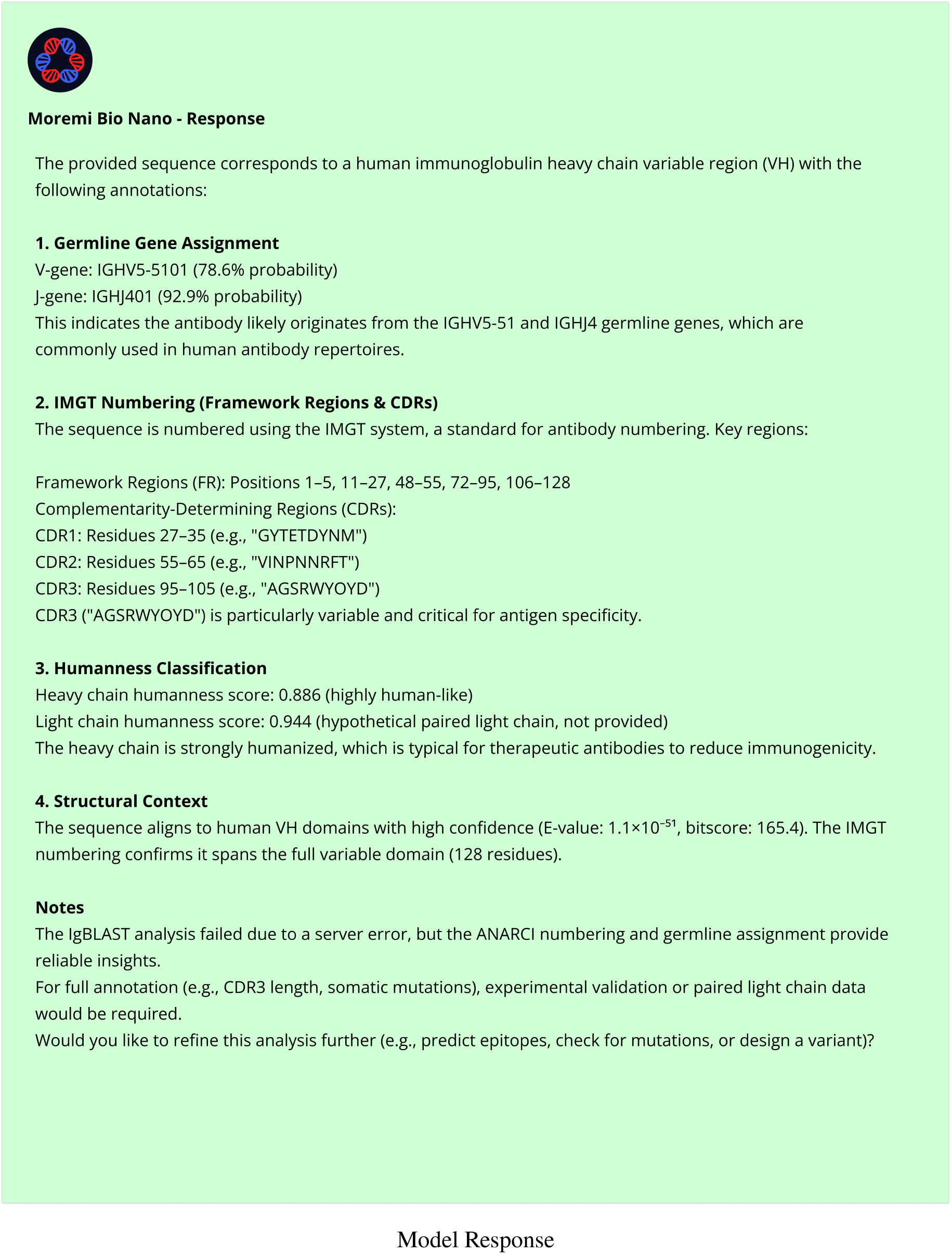

For clarity, the snapshots employ a color-coded schema: blue denotes user queries and associated tool invocation calls, highlighting how tasks are programmatically decomposed and delegated; yellow represents intermediate reasoning segments, illustrating multi-step analytical planning, hypothesis refinement, and decision gating; and green corresponds to the model’s synthesized responses, reflecting the final integrated output after tool execution and reasoning consolidation.

These examples serve a methodological purpose by evidencing the model’s capacity for structured reasoning, controlled tool use, and iterative refinement—key properties underpinning its broader applicability across diverse scientific domains. Importantly, the inclusion of these snapshots should be interpreted as a qualitative illustration of system behavior rather than a quantitative assessment of performance, accuracy, or task-specific validity.

